# A Simplified Method for Comprehensive Capture of the *Staphylococcus aureus* Proteome

**DOI:** 10.1101/2024.08.07.607079

**Authors:** Emilee M. Mustor, Jessica Wohlfahrt, Jennifer Guergues, Stanley M. Stevens, Lindsey Neil Shaw

**Author notes:** Corresponding authors Dr. Lindsey Shaw, Director, Center for Antimicrobial Resistance, Professor & Associate Chair Department of Molecular Biosciences, University of South Florida, 4202 East Fowler Avenue, ISA2015, Tampa, FL 33620-5150, USA, Dr. Stanley Stevens Jr., Professor, Department of Molecular Biosciences, University of South Florida, 4202 East Fowler Avenue, ISA2015, Tampa, FL 33620-5150, USA. These authors contributed equally to this work (Emilee M. Mustor, Jessica Wohlfahrt).

## Abstract

*Staphylococcus aureus* is a major human pathogen causing myriad infections in both community and healthcare settings. Although well studied, a comprehensive exploration of its dynamic and adaptive proteome is still somewhat lacking. Herein, we employed streamlined liquid- and gas-phase fractionation with PASEF analysis on a TIMS-TOF instrument to expand coverage and explore the *S. aureus* dark proteome. In so doing, we captured the most comprehensive *S. aureus* proteome to date, totaling 2,231 proteins (85.6% coverage), using a significantly simplified process that demonstrated high reproducibility with minimal input material. We then showcase application of this library for differential expression profiling by investigating temporal dynamics of the *S. aureus* proteome. This revealed alterations in metabolic processes, ATP production, RNA processing, and stress-response proteins as cultures progressed to stationary growth. Notably, a significant portion of the library (94%) and proteome (80.5%) was identified by this single-shot, DIA-based analysis. Overall, our study shines new light on the hidden *S. aureus* proteome, generating a valuable new resource to facilitate further study of this dangerous pathogen.

## Background & Summary

*Staphylococcus aureus* is a serious public health threat, causing infection in almost every ecological niche of the human body. It does this through the coordinated production of an arsenal of virulence factors, each of which is subject to extensive regulation in response to environmental cues. Indeed, due to their single cell nature, protein dynamics within bacteria such as *S. aureus* are fluid, facilitating not only their ability for adaptation but moreover, to engender infection. This ever-evolving nature of the bacterial proteome is driven by protein modifications, protein turnover, protein-protein interactions, and the need to adapt temporally in response to stimuli. As such, proteome resolution is critical for understanding the intricacies of cellular processes and the underlying dynamics that influence the pathogenesis of this bacterium. Thus, there is a clear need for technological advancements that provide an efficient, robust, sensitive, and reproducible way to analyze convoluted and dynamic bacterial proteomes. Moreover, the fundamental existence of the dark proteome, defined as a subset of proteins for a given organism that have historically gone undetected experimentally, also underlines the need for new techniques and approaches [1, 2]. The significance of comprehensive bacterial proteome capture and the possibility of bringing the dark *S. aureus* proteome further towards the light, is that resolution of these proteins could provide a deeper fundamental understanding to the biology and disease-causing abilities of this pathogen.

The field of mass spectrometry (MS)-based proteomic research has significantly evolved over time, with ion mobility (IM) separation combined with MS emerging as a powerful approach for the comprehensive characterization of proteomes [3, 4]. One specific area of development has utilized trapped ion mobility spectrometry (TIMS) combined with parallel accumulation serial fragmentation (PASEF) on a QTOF instrument (timsTOF), which has been shown to boost confidence in positive identifications and overall proteome coverage [5-9]. As co-fragmentation of peptides within MS/MS acquisition is a complicating factor for in-depth proteome coverage, TIMS can partially address this issue by providing an additional dimension of peptide separation based on IM, which is related to its collisional cross section (CCS) [5, 7]. We previously demonstrated that IM values measured using a TIMS-TOF instrument could be strategically implemented in the process of gas-phase fractionation of ions, providing enhanced peptide and protein identifications for deeper proteomic analysis. Specifically, we achieved a 19% and 30% increase in protein and peptide identification, respectively, when applying these methods to a human (HeLa) cell lysate [10]. Additionally, downstream DIA applications using the TIMS fractionated library allowed for high-resolution protein and peptide identification of the same sample material at even lower sample concentrations, as well as with shorter LC-MS analysis times [10].

In this study, we sought to adapt this new methodology to study the major human pathogen, *Staphylococcus aureus*. To date, the most comprehensive proteome for this organism came from a study by Michalik and colleagues, who generated an *S. aureus* DIA peptide ion-library reflecting an estimated proteome coverage of 72% [11]. Achieving this result required multiple strains of *S. aureus*, alongside varied environmental conditions and growth phases for library generation [11]. Given recent technological advancements in the field of mass spectrometry-based proteomics, we aimed to build upon the study by Michalik et al., shrinking the percentage of the *S. aureus* proteome that remains undetected, whilst at the same time increasing experimental throughput and limiting sample consumption. To do this, we used a single strain of *S. aureus*, at one growth time point without any environmental variability. We extended our methodology to include both TIMS fractionation as well as a simplified liquid-phase fractionation to further improve proteomic depth of coverage with minimal hands-on experimental time.

After generation of our comprehensive DDA-PASEF *S. aureus* spectral library, we then showcase its application for differential expression profiling by investigating temporal dynamics of the *S. aureus* proteome in a DIA single-shot global analysis.

## Methods

### Library Generation

#### Cell Culture

Five mL of TSB were inoculated in triplicate with an isolated colony of *S. aureus* USA300 LAC and incubated at 37°C overnight for 18 hours. Bacterial cultures were then synchronized to the same growth phase by the addition of 50 µL of the overnight culture to 5 mL of fresh TSB, followed by incubation at 37°C for 3 hours. Following this, to obtain an OD of 0.05, the required volume of the 3h culture was calculated and then added to 50 mL of fresh TSB and grown for 8h at 37°C. Cultures were then centrifuged and cell pellets washed three times with phosphate buffered saline (PBS) solution.

Meanwhile, to isolate secreted proteins, 10% trichloroacetic acid was added to the collected supernatants, and incubated at 4°C for 24 hours, followed by centrifugation to collect secreted protein precipitates. With regards to cell pellets, to facilitate lysis and protein extraction, samples were first resuspended in PBS containing 100 µg/mL lysostaphin, 100 U/mL DNAsel (Thermo Fisher), 25 U/mL RNAsel (Thermo Fisher), and 2x EDTA free protease inhibitor cocktail (Pierce, Thermo Fisher) before being incubated for 30 minutes at 37°C. Resuspended cell samples were transferred to a new vessel containing silica beads and subjected to 5 rounds of bead beating for 45 seconds each, with 1 minute rest intervals in between. Subsequent centrifugation at 17,000 G for 10 minutes fractionated cell samples further into soluble (supernatant) and insoluble (pellet) fractions. The insoluble fraction was washed with PBS and centrifuged at 17,000 G for 5 minutes.

#### Sample Processing

All samples were then prepared for mass spectrometry according to the Protifi S-trap protocol [12]. Specifically, samples were resuspended in cell lysis buffer with a final concentration of 5% (w/v) SDS, 50mM TEAB pH 7.55. Protein samples for each fraction were reduced by adding 20mM DTT and incubating at 95°C for 10 minutes. Next, to remove any non-solubilized proteins, samples were centrifuged at 17,000 G for 10 minutes and supernatants collected. All samples were then quantified using the Pierce 660nm assay with the ionic detergent compatibility reagent (Thermo Fisher Scientific); and standardized to 50 µg. Protein alkylation was performed via the addition of iodoacetamide to a final concentration of 40mM, followed by incubation at room temperature away from light for 30 minutes. The reaction was quenched with 12% phosphoric acid before adding 7 times the sample volume of the S-trap buffer containing 90% methanol and 100mM TEAB. Samples were applied to the S-trap column and centrifuged at 4,000 G for 30 seconds followed by three washes using S-trap buffer. Trypsin/Lys-C was added in a ratio of 1:10 and samples were digested overnight at 37°C. Following incubation, peptides were eluted via a series of three centrifugation steps, the first using 50mM TEAB, followed by 50mM TEAB with 0.2% formic acid, and finally, 50% acetonitrile. Eluted peptides were placed in a vacuum centrifuge until dry. Dried peptides were resuspended in 0.1% formic acid in water and desalted using C18 columns (Waters) according to manufacturer’s instructions. Peptides were then dried again using vacuum centrifugation.

#### iST peptide fractionation

Peptides for empirical library generation were fractionated using the iST peptide fractionation add-on (PreOmics GmbH). *S. aureus* peptides were resuspended in iST Lyse buffer. The insoluble and soluble fractions were pooled, while the secreted fraction was kept separate. Resuspended peptides were added to an iST cartridge and washed with iST Wash buffers 1 and 2, each followed by a centrifugation step of 3,800 G for 2 minutes. Peptides were eluted into 3 separate tubes during 3 rounds of centrifugation at 1,000 G for 1 minute with the appropriate iST Fraction Elution buffer. Once dried in a vacuum centrifuge, peptides were resuspended to a concentration of 2 µg/µL with 0.1% formic acid in water for library generation.

#### UHPLC gradient profile

To characterize the *S. aureus* proteome, a nanoElute UHPLC system (Bruker) with reversed-phase C18 column (Aurora series generation 3, 25 cm x 75 µm i.d., 16 µm C18, IonOpticks) heated to 50°C was used to separate samples in conjunction with a TIMS-TOF mass spectrometer (timsTOF Pro, Bruker) for in-line LC-MS analysis. A gradient of 2-25% B, with mobile phase A being 0.1% formic acid in LC/MS-grade water and B 0.1% formic acid in acetonitrile, was used for peptide elution. Following this step, the gradient was ramped up to 37-80% B to further elute remaining hydrophobic peptides and prepare the column for subsequent samples.

#### LC-MS/MS analysis and data processing

In order to generate the empirical library, a gradient length of 120 min (160 min total gradient time with column wash) was implemented on all three iST peptide fractions in triplicate using DDA-PASEF analysis and two IM fractions, 0.8-1.05 1/K_0_ [V•s/cm^2^] and 1.0-1.25 1/K_0_ [V•s/cm^2^] as previously described [10]. A non-fractionated sample analyzed by DDA within the IM range of 0.7-1.4 1/K_0_ [V•s/cm^2^] was also acquired to allow for proper alignment of IM and retention time (RT) using EasyPQP and to identify the remaining limited peptides within the outer edges of the IM range used for DIA acquisition. DDA-PASEF method settings included a range of 100-1700 m/z, ramp time of 100 ms, a ramp rate of 9.43 Hz, and an active exclusion list that releases after 0.4 minutes. Additionally, the total cycle time was set at an estimated 1.17 seconds with a target intensity of 15,000 and an intensity threshold of 2,500. After acquisition, data were compiled into a library file (.TSV) using FragPipe (v.19.1) with the DIA_SpecLib_Quant workflow. The following FragPipe tool versions were utilized: MSFragger (v.3.7), IonQuant (v.1.8.10), Philosopher (v.4.8.1) EasyPQP (v.0.1.36). FragPipe with the LFQ-MBR workflow and MaxLFQ min ions set at 1 was also used to search these files separately based on replicate number (n=3) to assess overall quantifiable protein and peptide coverage and reproducibility of the experimental fractionation approach.

### Differential expression profiling

#### Sample preparation

Samples were prepared as detailed above without iST fractionation, however cultures were grown to 3, 6, and 16h, to facilitate capture of mid exponential phase, entry into stationary phase, and end stationary phase, respectively.

#### LC-MS/MS analysis and data processing

DIA-PASEF analysis on the TIMS-TOF instrument was used to characterize these different growth phases over a 90-minute gradient. DIA-PASEF was acquired over one range of 0.7-1.4 1/K_0_ [V•s/cm^2^], with a 47-mass step polygon window, each with a mass width of 25 Da and no overlap. An m/z range of 250-1425 was used, with an estimated cycle time of 1.48 seconds. These data were searched in DIA-NN (v.1.8.1) using the previously generated empirical spectral library with mass accuracy and MS1 accuracy set to 15 ppm. Isotopologues were used, and MBR was enabled. Protein inference was set to genes, single-pass mode was selected as the neural network classifier, robust LC (high precision) was set for the quantification strategy, cross-run normalization was set to RT-dependent, and smart profiling was set for library generation.

#### Data filtering and statistical analysis

Cellular and secreted fractions at the 3 time points were then analyzed in Perseus (v.2.0.11.0) separately, to assess dataset quality and perform further data processing.

Samples were analyzed with at least 4 out of 4 valid values in at least one time point group detected. The remaining missing values were replaced using the imputation function with a width of 0.3 and downshift of 1.9 for cellular fractions and a width of 0.5 and a downshift of 1.6 for secreted fractions to fit the imputed values into the lower abundance area of the Gaussian curve. A one-way ANOVA was performed followed by a post hoc Tukey’s HSD test (FDR<0.05) to determine significance of differential expression when comparing the proteomes at different time points (16vs3h, 16vs6h and 6vs3h). In the **Supplemental** Excel, proteins with purple font indicate that at least one of the two groups in that comparison have 2 or fewer originally measured values only (before imputation, shown further right on the excel sheet with columns labeled “original”). The values in red cells indicate that imputation occurred on all 4/4 reps and no measurement was provided through the instrumentation.

## Data Records

All mass spectrometry data have been deposited to the ProteomeXchange Consortium via the PRIDE [13] partner repository with the dataset identifier PXD053994.

## Technical Validation

The initial part of the workflow utilized in this study was deep proteomic analysis of *S. aureus* using DDA-PASEF to generate a comprehensive spectral library. Technical validation was achieved in both peptide and protein identification confidence as well as experimental reproducibility of generating the empirical library. MSFragger was used to search the DDA-PASEF data within the default DIA_SpecLib_Quant workflow of FragPipe. Following this search, MSBooster, Percolator, and ProteinProphet were utilized in FragPipe to further improve confidence in peptide and related protein identifications for subsequent library creation. Output from these validation steps were input into EasyPQP, resulting in a final, high-confidence library file (.TSV) consisting of only peptides that had passed a 1% FDR cut off for proteins, peptides, and peptide spectral matches. This library file was utilized in subsequent DIA-NN searches of DIA-based analysis of *S. aureus* in the context of a comprehensive, differential expression profiling experiment. For this experiment, temporal proteome changes during cell culture were characterized, with an FDR cutoff at the precursor (global and run-specific) and protein level (global) of 1.0% within the DIA-NN analysis. The protein groups matrix, with an additional 5% run-specific FDR at the protein level, was utilized for further filtering and statistical analysis in Perseus. The dataset was filtered to include only consistently detected proteins and differential expression significance established at FDR<0.05.

To validate quantitative precision of the empirical library generation, technical replicates of the entire workflow consisting of sample processing, fractionation and LC-MS analysis were performed. In terms of quantitative precision of just the cellular fraction, 1,942 proteins out of the 2,074 proteins detected in all three replicates of the cellular fraction recorded a coefficient of variation (CV) of less than 30% (**Figure 1A&B**), with an average and median CV of 14.7% and 8.4%, respectively. Secreted fractions were analyzed separately from the cellular fractions to facilitate more comprehensive characterization of these secreted proteins, which are found in lower abundances within the cell and inherently generate a lower peptide coverage. In general, the secreted fractions exhibited higher variation, with an average CV of 53% and median of 26%. While these fractions showed lower quantitative precision compared to the cellular fraction, they did allow deeper coverage at the peptide level in order to significantly enhance the total library size (**Figure 1B**). Related to the differential expression profiling experiment, percent CV calculations demonstrated lower variation within each timepoint for DIA-based proteomic analysis of the cellular fraction samples using the empirical library, with an overall mean and median CV of 17% and 14%, respectively. For DIA-based proteomic analysis of the secreted fraction, the mean CV for all time points together was 36%, while the median was 27%, which highlights the expected higher biological and experimental variability associated with these samples.

**Figure 1.**
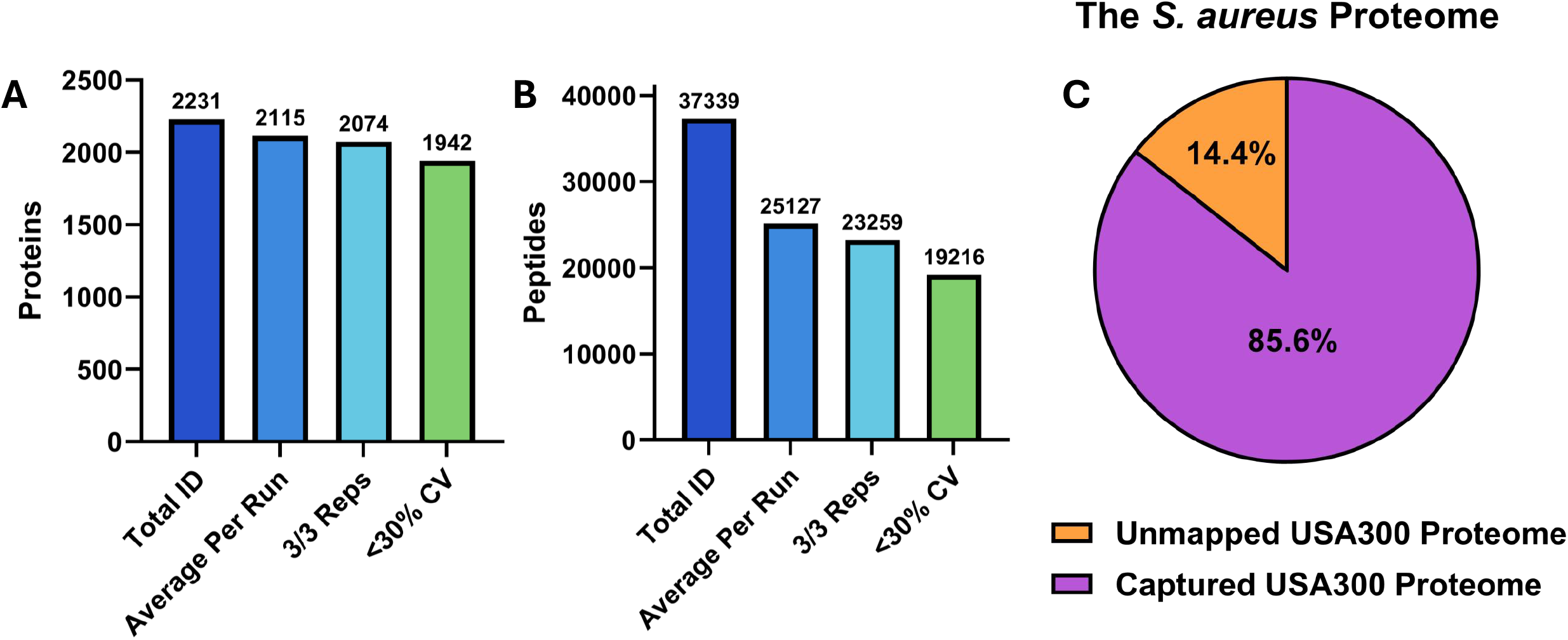
Comprehensive Capture of the *S. aureus* Proteome. Number of proteins (**A**) and peptides (**B**) identified when searching all samples to compile a *S. aureus* empirical spectral library. This was achieved using a 120-min gradient and two IM fractions: 0.8-1.05 1/K_0_ and 1.0-1.25 1/K_0_. (**C**) Percentage of the *S. aureus* USA300 proteome identified in our library.

## Usage Notes

### Library generation

Overall, our approach facilitated the capture of, to the best of our knowledge, the highest number of experimentally detected unique *S. aureus* proteins to date. Across all samples, biological replicates as well as iST peptide cellular and secreted fractions, we recorded a total of 2,231 proteins (**Table S1** and **Figure 1A**) and 37,339 unique peptides (**Table S2** and **Figure 1B)**. To assess the reproducibility of the entire approach, which includes variability of both the liquid- and gas-fractionation based on individual replicates, we separated individual replicates and performed LFQ analysis for the cellular fraction (n=3). An average of 2,115 ± 10 quantifiable proteins based on measured MaxLFQ intensities were identified per run. Collectively, these data encompassed 85.6% of the *S. aureus* USA300 LAC proteome (**Figure 1C**) and demonstrated the high reproducibility of our approach even with the addition of both gas- and liquid-phase fractionation. By comparison, Michalik and coworkers identified 2,057 *S. aureus* proteins (72% coverage) corresponding to 25,664 peptides [11]. Their library was achieved using multiple growth phases, multiple strains of *S. aureus* cultured in two types of media, and experimental variations that included manipulated iron concentrations [11]. In total, that study performed 152 proteome measurements to compile their DDA spectral library. Contrastingly, our methodology used 3 liquid fractions and 2 gas-phase (TIMS ranges), totaling 6 injections per replicate with 200ng digest loaded per injection. Using both cellular and secreted fractions, this corresponds to approximately 2.4 µg of total material with 12 LC-MS analyses per replicate.

The data compiled in this library provides the most comprehensive proteome of *S. aureus* to date and are publicly available to serve as a resource for subsequent studies. We envision that this dataset can build upon previously established studies and related online resources such as that presented by Depke et. al [14] to facilitate new and novel studies exploring the host-pathogen interface.

### Differential expression profiling

After generation of our comprehensive DDA-PASEF spectral library, it was then used for DIA single-shot global analysis in the context of a differential expression profiling experiment. Accordingly, we investigated the temporal dynamics of the *S. aureus* proteome during different growth phases to facilitate capture of mid exponential phase (3h), entry into stationary phase (6h), and end stationary phase (16h), respectively. Upon analysis of the various time point samples, we observed high reproducibility demonstrated via principal component analysis (PCA) (**Figure 2**). Cellular and secreted fractions revealed clustering of samples within individual timepoints, except for one replicate from the 16h secreted dataset (**Figure 2A&B**). Importantly, PCA also demonstrated distinct separation of the time-point groups, indicating robust proteome changes underlying the different growth phases. Given that we recorded high sample reproducibility, our next focus was to discern differentially expressed proteins to distinguish how the proteome changed over time. We identified a total of 2,101 unique proteins across all replicates, time points, and cell fractions. While analysis of the secreted fraction for each time point provided us with valuable information (**Table S3**), for brevity our focus was on the cellular fraction, where we identified 1988 ± 14 proteins per run across all time points. In comparison to Michalik et al., whose study identified an average of 1432.5 proteins per DIA analysis, this is an approximate 40% improvement in single-shot proteome coverage [11]. Following analysis of the cellular fraction, we identified 708, 1178, and 736 proteins with significantly altered production in the 16vs6, 16vs3, and 6vs3 timepoint comparisons, respectively (**Table S4**). ShinyGO enrichment analysis using the corresponding genes presented a holistic view of how the cellular proteome changed over time (**Figure 3A-C**). In accordance with existing literature, we observed significant changes in metabolic and biosynthesis processes, catalytic activity as well as other cellular pathways such as metal and ion binding in the 16-versus 3-hour datasets (**Figure 3A&B**). The metabolic profiles of *S. aureus* cells in exponential phase are known to drastically contrast those in stationary phase due to a transition in the origin of ATP production [15]. Hence, this reasoning could also account for why we observed almost double the number of proteins that exhibited significant changes in the 16hvs3h analysis compared to the 16hvs6h and 6hvs3h evaluations. However, when looking at proteome changes between 16h and 6h, this analysis presented unique changes in comparison to other timepoints, indicative of the transition from exponential to stationary growth. Once bacterial cells enter stationary phase, they undergo several morphological and physiological changes to not only account for but combat stress, such as nutrient depletion, the presence of toxic metabolic by-products, as well as alterations in pH and osmolarity [16, 17]. For example, we observed the largest fold enrichment in chaperone proteins known to maintain proper protein folding following oxidative damage [18]. Furthermore, we recorded an increase in manganese utilization moving into stationary phase (**Figure3A**). This represents an additional avenue to protect against oxidative stress as this ion is a critical co-factor for superoxide dismutase, a neutralizer of reactive oxygen species. Lastly, in this comparison is the observed alteration of tRNA processing (**Figure 3A**). Indeed, mRNA and tRNA accumulation has been reported as a bacterial defense response when faced with nutrient starvation [16, 19]. Overall, the empirical library-based methodology was well-suited for high-resolution and comprehensive proteome capture, especially for smaller-scale proteomes such as that of *S. aureus*.

**Figure 2.**
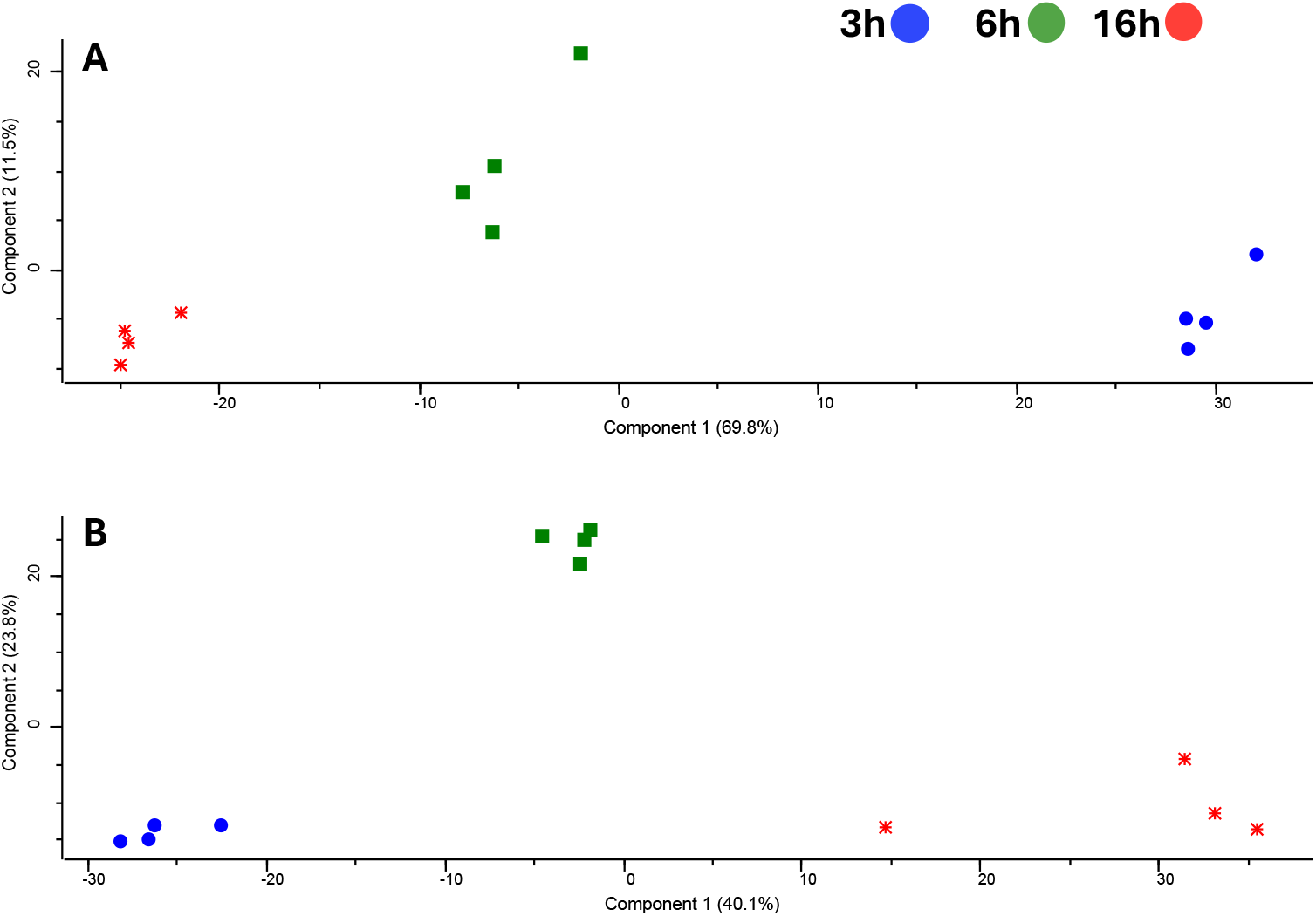
Principle component analysis (PCA) of *S. aureus* proteomes sampled from three different growth phases. PCA plots highlight sample reproducibility shown by tight clustering of replicates for (**A**) cellular and (**B**) secreted fractions at 3 hours (blue), 6 hours (green) and 16 hours (red).

**Figure 3.**
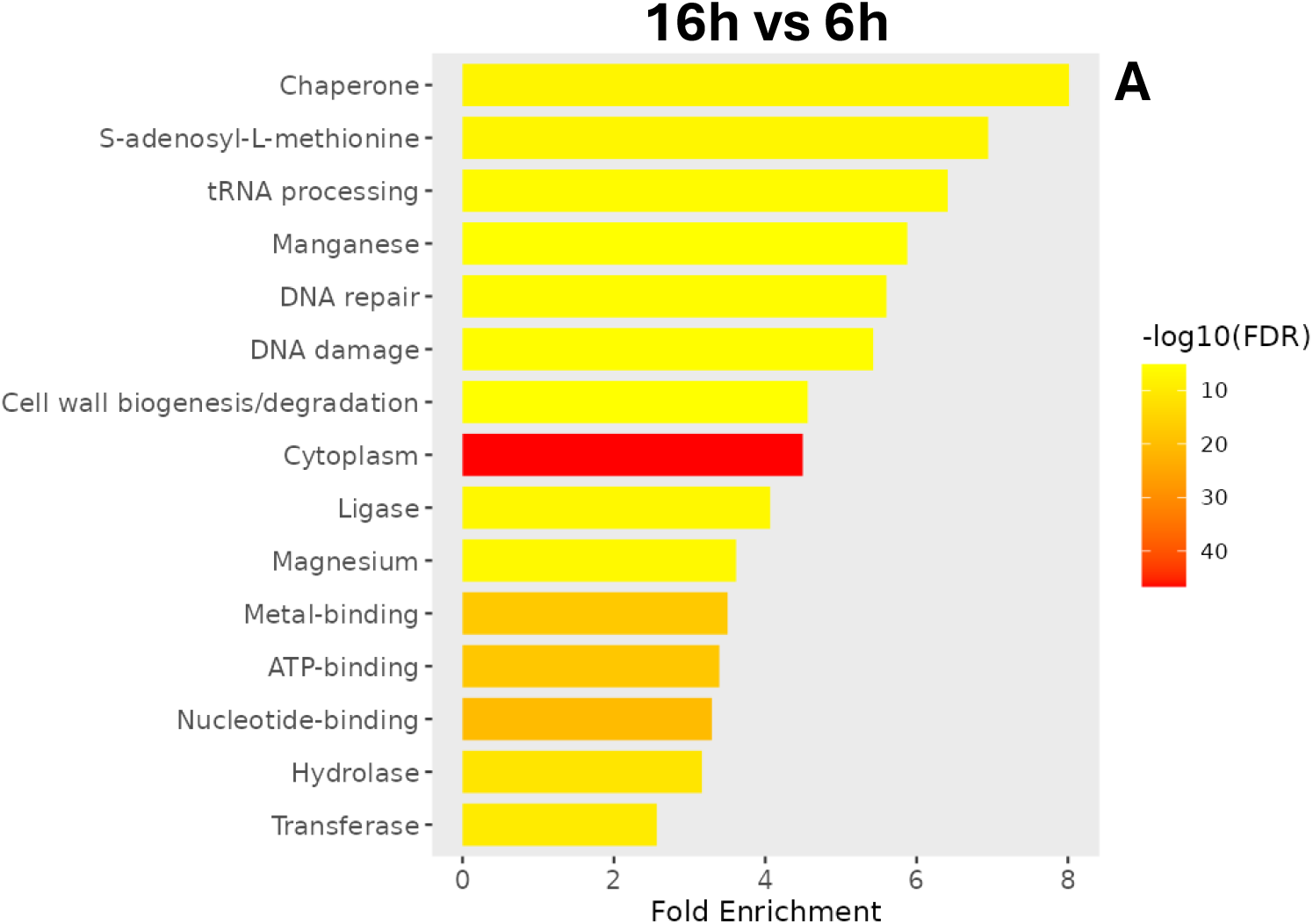
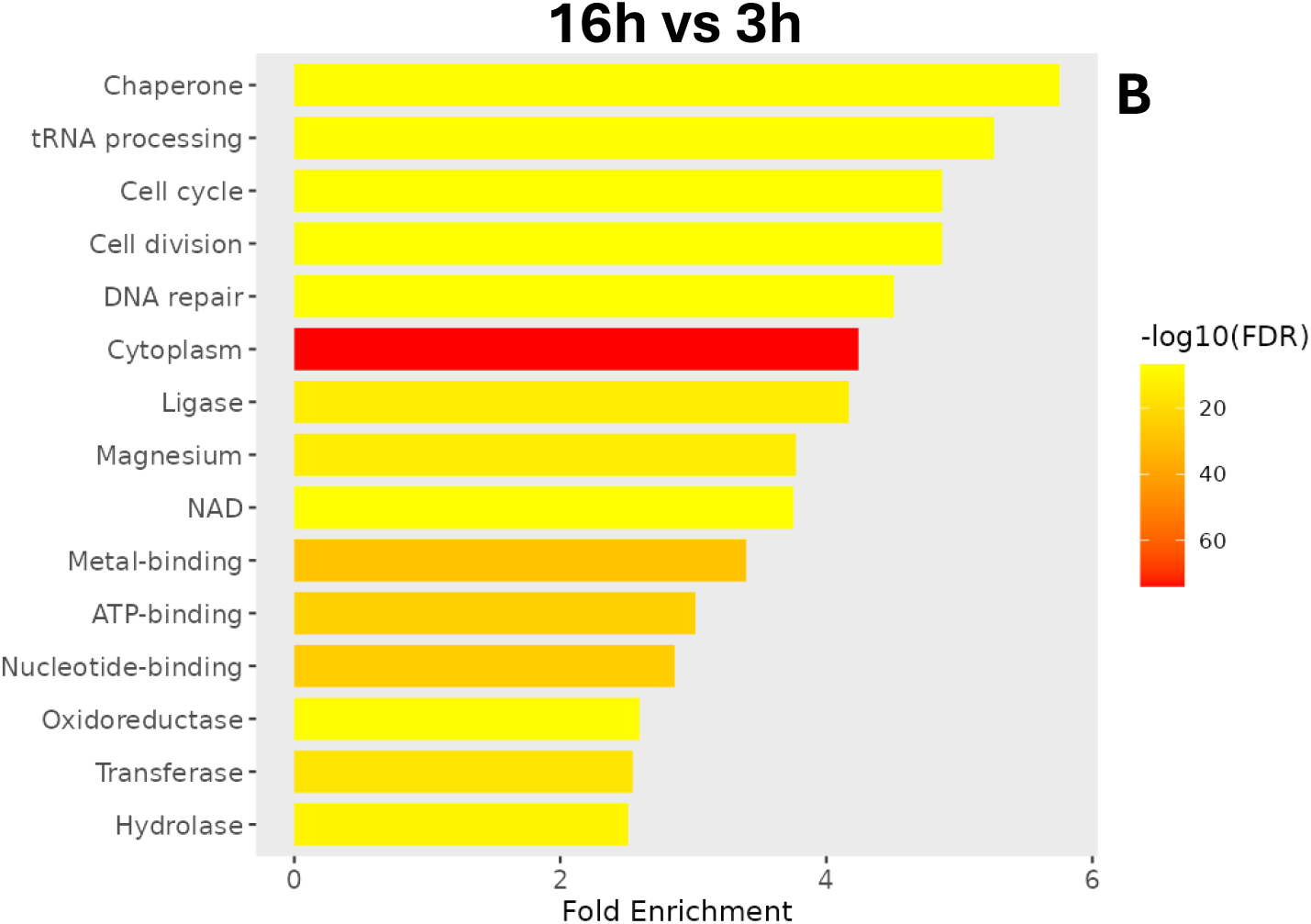
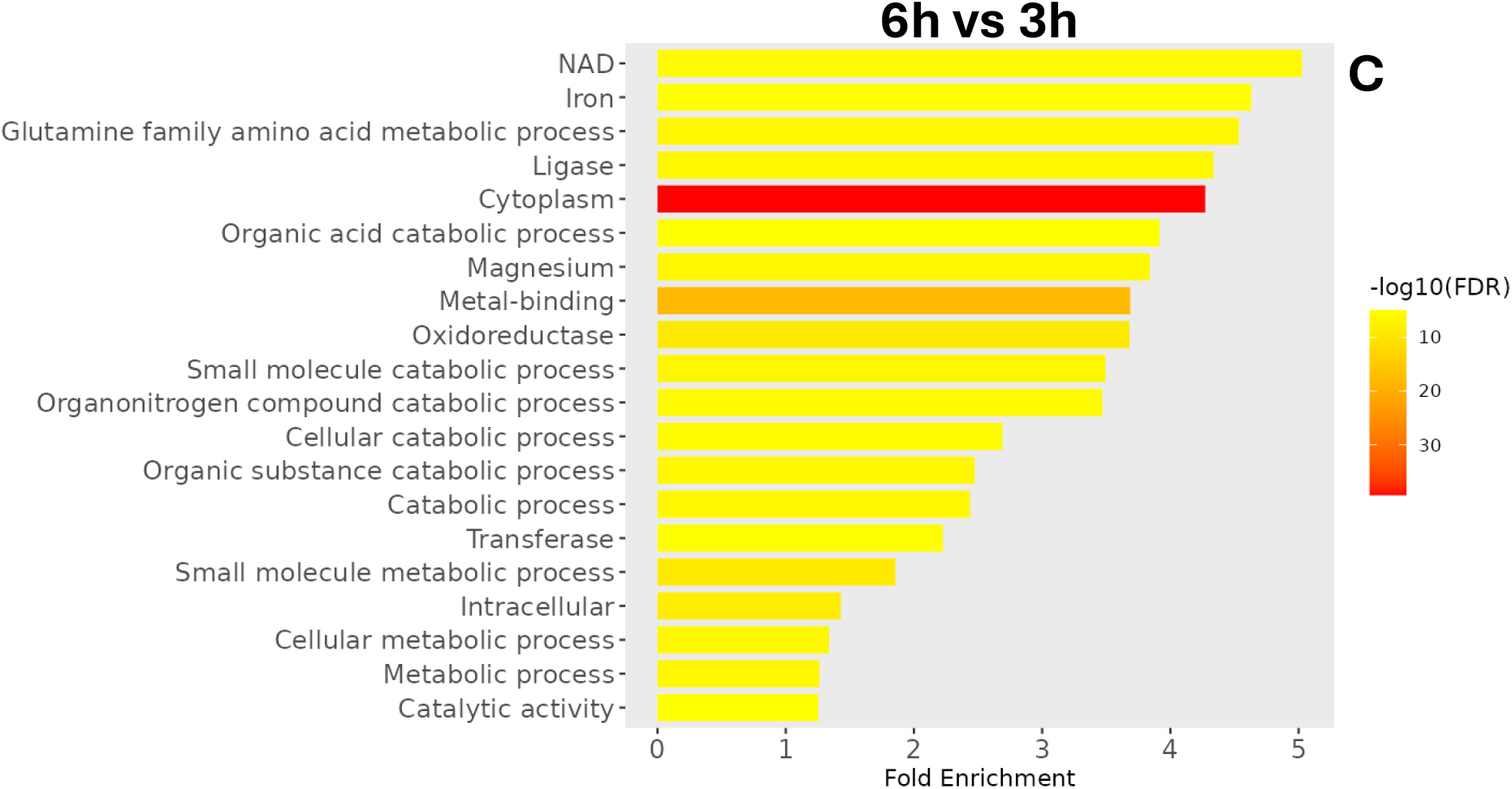
Ontological enrichment analysis of proteins with differential expression profiles during distinct growth phases. Shown are the top 20 pathways retrieved from ShinyGo for the comparison of (**A**) 16 vs 6 hours, (**B**) 16 vs 3 hours, and (**C**) 6 vs 3 hours. Pathway titles correspond to the annotated keywords in the Uniprot database. Fold enrichment and false discovery rates (FDR) refer to the input gene list to determine overrepresentation.

This present study is the first to demonstrate the utility of a streamlined liquid- and gas-phase (ion mobility) fractionation approach in combination with PASEF analysis on a TIMS-TOF instrument to further understand biological systems in prokaryotic organisms. Consequently, we generated the most comprehensive *Staphylococcus aureus* proteome to date. Given the sensitivity and depth of coverage attainable by the TIMS-TOF instrument, a single-shot analysis of *S. aureus* lysate digest using DDA-PASEF acquisition alone can identify a significant portion of its proteome using a 90 min UHPLC gradient (∼70%), which was improved to over 85% (i.e., 2,231 total proteins) in the empirical library generated herein. Although protein identification improvement through our gas- and liquid-phase fractionation of the *S. aureus* proteome is modest compared to single-shot analysis, we note additional gains in protein identification to achieve extended coverage within the dark proteome will likely require a significant amount of experimental and instrument time [2]. Also, given the speed of data acquisition on the TIMS-TOF instrument, the LC-MS analysis time could be significantly reduced without much compromise in depth of coverage for this particular proteome. However, we envision the library and raw data available through PRIDE from this study will be a valuable resource to the community given the high degree of peptide separation performed along with CCS values that can be derived from recorded IM values (e.g., future DIA or PRM applications). While our study was limited to only common variable modifications such as methionine oxidation and consideration of fully tryptic peptides, leveraging this data, additional post-translational modifications could be investigated. This includes more open searches of enzyme specificity to significantly increase the peptide library size to account for proteolytic activity of endogenous proteases that are important to the cellular stasis and pathogenesis of *S. aureus* [20, 21].

Using this comprehensive empirical library, we observed the discernible advantages of minimized hands-on experimental time and sample consumption for downstream DIA analysis of the different growth phases of *S. aureus*. This application demonstrated a marked improvement in single-shot proteome coverage with DIA analysis (∼40% more proteins identified) of *S. aureus* compared to previously published studies [13]. Although not presented in this study given the focus on generation and implementation of an experimentally derived library, a library-free approach could also be used in combination with the empirical library to potentially complement and enhance protein identification from DIA analysis. Collectively, our studies have generated a valuable protein catalogue for *S. aureus*, which has been effectively utilized to provide detailed molecular insight into temporal proteome dynamics. This study further demonstrates the recent advancements in the field of mass spectrometry for improved proteome coverage, presenting an invaluable avenue to understanding the biology of drug-resistant pathogens.

## Code Availability

All version numbers of the software utilized are listed in the methods section. No custom code was utilized in this study.

## Acknowledgements

This study was supported by grants AI124458 and AI157506 (both to L.N.S.) from the National Institute of Allergy and Infectious Diseases.

## Author contributions

LNS and SMS designed the project, EM performed cell culture experiments and collected samples, EM and JW processed samples and generated figures, JW and JG performed LC-MS, JG performed data analysis. All authors contributed to the writing of the manuscript.

